# Estimation the porosity of the valves of centric diatom *Minidiscus vodyanitskiyi* Lyakh & Bedoshvili using SEM images

**DOI:** 10.1101/523258

**Authors:** Anton M. Lyakh, Yekaterina D. Bedoshvili, Olga V. Shikhat

## Abstract

The diatoms interact with the outer environment through the siliceous walls of a frustule. Because of that the surface area of the frustule determines the ability of diatoms to absorb and excrete material resources. Such as unicellular organisms exchange matter only through the pores in their cell wall, to find relationships between characteristics of material fluxes and surface area of microorganism cover that is penetrable for substance, it is necessary to estimate the total surface area of pores or a porosity – relative area of pores perforated frustule. In the paper we describe a method of estimating the porosity of a diatom valve using SEM images. The method is tested on SEM images of the valves of centric diatom *Minidiscus vodyanitskiyi* recently described from the Sea of Azov. The results show that the valves porosity is less than 5 % of the total valves area. This value is consistent with the relative perforation of land plants leaves, which is less than 3%. We hypothesize that such value of diatom valves porosity is usual for many other diatom species. To verify this hypothesis additional researches are necessary.

## Introduction

The diatoms, as many other unicellular organisms, interact with the outer environment through the siliceous walls of a frustule. The architecture of frustule surface determines the ability of a diatom to exchange the nutrients and the products of vital function [Bukhtiyarova 2009; Hale & Mitchell 2001; Pahlow et al. 1997; Pickett-Heaps et al. 1990]. The area of a cell cover outlines individual contact space, where “physical, maximum energy and informative interaction, high interchange of substances are accomplished between individual and environment” [Bukhtiyarova 2013]. The frustule surface area-to-volume ratio characterizes the intensity of those processes [Hein et al. 1995; Pahlow et al. 1997]. However when researchers estimate the surface area of microalgae, they do not take into account that unicellular organisms exchange matter only through the pores in their cell wall. The other parts of a cell wall are impenetrable for material fluxes. Therefore if we examine the fluxes of material through the cell wall relying on the area of pores rather than the total surface area, we can improve existing relationships between characteristics of material fluxes and organism morphometry or discover new ones.

*The porosity* of the microalgae cover is the relative surface area perforated with pores and potentially penetrable for fluxes of matter.

In the paper we estimate the porosity of valves of centric diatom *Minidiscus vodyanitskiyi* (Fig. 1–3) recently described from the Sea of Azov [Lyakh & Bedoshvili 2018]. Their relatively high abundance allow collecting a set of SEM images where valve areolae and pores that perforated areolae cribra are well distinguished. The valves of those diatoms have tiny sizes, because of that they are covered by relatively small number of locular areolae, that helped us to estimate their porosity using image processing technique and manually selecting areolae on SEM images.

**Figures 1–3.**
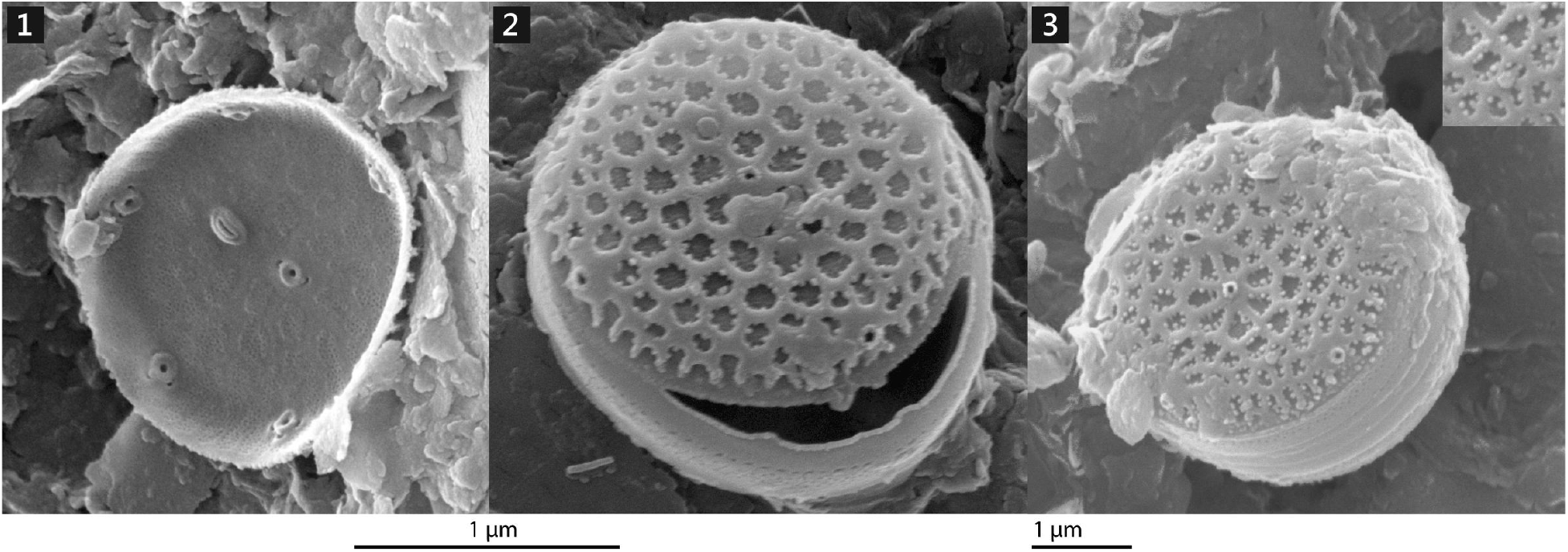
Inner (1) and outer views (2–3) of the valves and frustules of *M. vodyanitskiyi*. In the top-right corner of fig. 3 there is a zoomed part of a valve with siliceous growth on areolae ribs.

## Material and Methods

Study area, sampling and microscopic technique, morphology of *M. vodyanitskiy* frustule and valve are thorough described in the previous paper [Lyakh & Bedoshvili 2018].

Because the most of the studied diatoms were lied on valve, we can collect statistically significant data only about porosity of valves.

To find porosity of valves, we estimated the total area of all areolae on every valve; found the number and area of cribral pores, that perforate cribra of every areola; calculated cribral pores total area, and compared it with the surface area of a valve.

To fulfill every step, we used: GIMP image editor (gimp.org) to select areolae on SEM images; Inkscape vector editor (inkscape.org) to outline the valve border and regions of a valve surface covered by mineral granules; ImageJ image processing software (imagej.nih.gov/ij) to estimate the average diameter and distance between cribral pores; and an auxiliary software developed by the first author to estimate the valves porosity. The details of every step are described below.

*Step 1. Find the elliptical distortion of a valve*. The studied diatoms have circular valves [Lyakh & Bedoshvili 2018]. However on SEM images some valves have the shape of an ellipse which is sometimes rotated, because those valves do not lie in the image plane (Fig. 1–3). The images of such valves are elliptically distorted. The ratio between minor and major diameters of the ellipse, that outlines valves margin, determines the coefficient of distortion, *k*_ell_ = *D*_major_ / *D*_minor_. If we multiply the area of the ellipse on *k*_ell_, we obtain the area of the original circular valve. To find *k*_ell_, it is necessary to draw an ellipse along a valve border. It is known from the analytical geometry that any five points, every three of which are not lied on a straight line, define an unique ellipse. Therefore, to construct those ellipses, we loaded every image of a valve into Inkscape, put five points on valve margin using “Draw Bezier curves and straight lines” tool, and used “Ellipse by 5 points” Inkscape extension [Pernsteiner 2018], which turns a path with five control points into an ellipse passing through them (Fig. 4). The constructed ellipse consists of four control points such that pairs of opposite points define the main ellipse diameters. That allow easily finding those diameters (Fig. 4) and calculating coefficient of distortion. This technique allow restoring borders of any valves including particularly visible.
*Step 2. Selecting areolae on the valve image*. To select areolae on images of valves, we used two ImageJ segmentation algorithms: thresholding and watershed segmentation. Unfortunately they could not identify all areolae on every studied valve due to a low contrast between areolae and ribs, and the presence of inorganic material on a valve surface (Fig. 4–5). Because of that we manually selected other areolae in GIMP using “Fuzzy Select (Magic Wand)” tool. Selected areolae were painted in yellow or blue color depending on their position and connection with neighboring areolae. Areolae that are located on valve margin and touched each other were painted in blue color, all other in yellow color (fig. 5).
*Step 3. Restoring convex shape of areolae*. Because areolae on some valves are closed by filiform siliceous growths (Fig. 2–3, 6) [Lyakh & Bedoshvili 2018], the selected regions sometimes do not represent a real areolae shape. On the masked image that areolae have the shape of concave polygons (Fig. 7). To restore a real shape of areolae, we used original auxiliary software that makes areolae convex constructing a convex hull around them. That allow removing filiform growths from the areolae images (Fig. 8). The program makes convex only those regions (areolae) that are painted in yellow color and ignores others.
*Step 4. Identifying inorganic flakes on valve image*. The surface of many valves were covered by inorganic material (fig. 2–4). The area of that regions is excluded from total area of the valve. To highlight those regions in valve images, we manually outlined them by vector polygons in Inkscape and painted in purple color (fig. 5). This color indicates to our program that area of those regions should be extracted from the area of a valve.
*Step 5. Calculating the surface areas of a valve and areal of areolae*. The area of a valve surface, *A*_valve_, is equal to the area of a circle with the diameter *D* major, which is the major diameter of the bounding ellipse (fig. 4). The total areas of valve areolae, *A*_areolae_, and inorganic flakes, *A*_inorganic_, are determined as the total areas of colored spots on a valve image. The spots areas are equal to the number of pixels painted in the predefined colors. Yellow and blue colors correspond to areolae, pink color denotes inorganic flakes. The numbers of pixels of each color are counted by the developed program. These numbers were converted to square micrometers using the ratio of the length of a scale bar in micrometers and in pixels. To remove elliptic distortion, all calculated areas were multiplied to *k*_ell_. The area of a valve that is free from inorganic flakes, *A*_free-valve_, and relative area occupied by areolae, *A*_areolae-relative_, were calculated as follow:

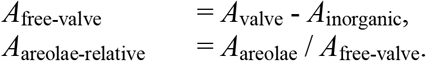
*Step 6. Estimating an approximate number and the total area of cribral pores*. Studied diatoms have locular areolae that are covered from the inner side by a siliceous plate named cribrum. The cribrum is equally perforated by small *cribral pores* (Fig. 9, 10) [Lyakh & Bedoshvili 2018], which facilitate matter exchange between protoplast and the environment. We assumed that all cribral pores were circles of equal diameters located on the same distances from each other. Using SEM images of the inner parts of valves, we estimated the average diameters of cribral pores, *D*_pore_, and the area of one pore, *A*_pore_ = π *D*_pore_^2^ / 4. To estimate the number of pores on a valve, *N*_pores_, we measure average distance between them, *D*_interpore_. SEM images show that cribral pores are distributed in a triangular grid (Fig. 9). We assumed that this grid is regular that allow replacing every pore with surround siliceous plate by a right hexagon with a pore in the middle (Fig. 10). The area of a hexagon is

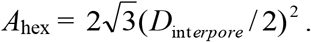 The number of hexagons, that can fit into the given area of areola, gives the approximate number of pores perforated that area. The task of finding the exact number of regular hexagons fit the give area does not have unambiguous solution. The approximate solution is to divided the given area on the area of one hexagon, and take the integer part of the resulting value. After that we can estimate the total area of all pores, *A*_all-pores_:

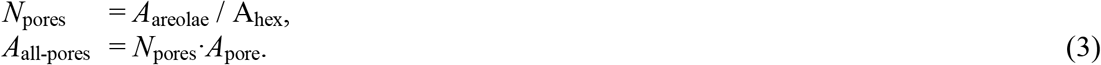
*Step 7. Estimating the porosity of a valve*. The porosity of a valve, *Y*_valve_, is the ratio of the total area of all cribral pores to the total area of a valve:

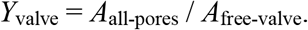

**Figures 4–10.**
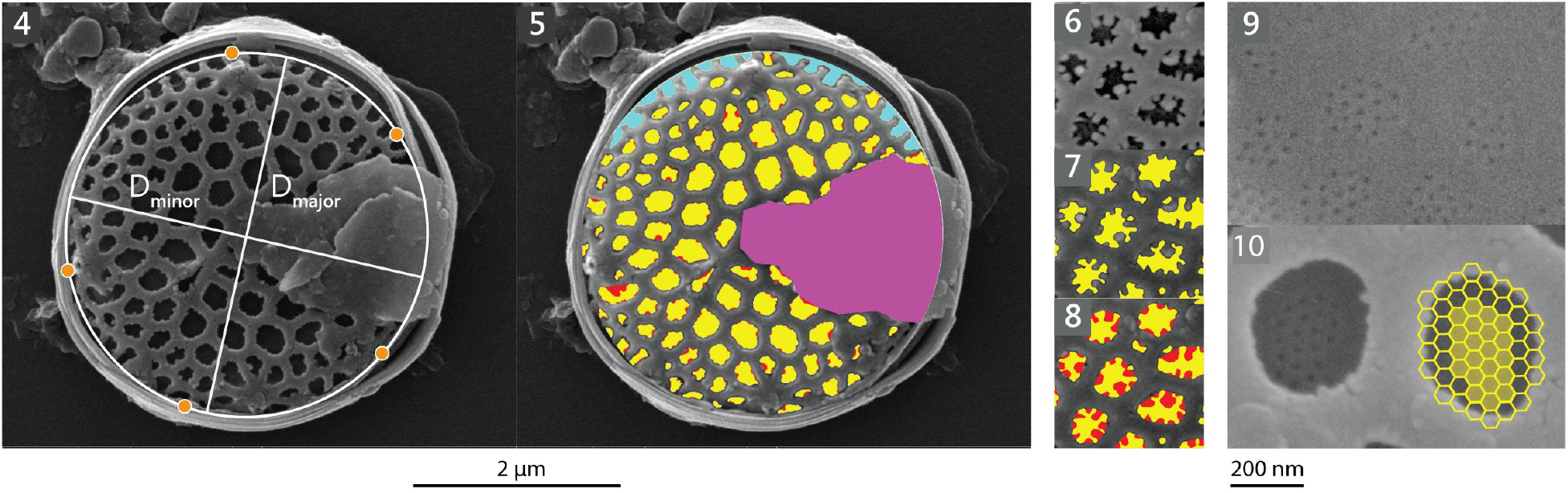
Steps used to measure valves porosity on SEM images. **4**. The bounded ellipse is constructed using five points, and ellipse main diameters were determined. **5**. Areolae (yellow and blue) and inorganic flakes (pink) were manually selected. **6–8**. The real convex shape of areolae covered by filiform siliceous growth was restored using convex hull algorithm. **9**. Average diameter and distance between cribral pores were measured. **10**. Cribral pores were represented by hexagons and the number of those hexagons was estimated.

## Results

The measured morphometric characteristics of *M. vodyanitskiyi* specimens and other centric diatom species are presented in Table 1.

**Table 1.** Comparison of morphometric data of three centric diatom species.

|  | *M. vodyanitskiyi*<br>(17 valves) | *T. eccentrica*<br>(1 valve)<br>[Losic et al. 2007] | *Coscinodiscus* sp.<br>(1 valve)<br>[Losic et al. 2007] |
| --- | --- | --- | --- |
| Valve diameter, $\mu\text{m}$ | 2.7–7.8 | 30–40 | – |
| Diameter of cribral pores, nm | 20 | $43 \pm 6$ | $45 \pm 9$ |
| Distance between centers of cribral pores, nm | 50 | – | $68 \pm 12$ |
| Relative area of areolae, % | $34 \pm 5$ | $37 \pm 3$ | $35 \pm 3$ |
| Porosity of an areola, % | $15^{1)}$ | $10 \pm 2.5$ | $7 \pm 1.5$ |
| Porosity of a valve, % | $5 \pm 0.7$ | $3.7^{2)}$ | $2.5^{2)}$ |

<sup>1)</sup> The calculated porosity of areolae is independent on the total areolae area; the real areolae porosity is smaller.
<sup>2)</sup> These values are calculated as the product of the porosity of areolae on the relative area of areolae.

The calculated number and total area of cribral pores is bigger than real ones, because pores do not perforate full cribra area but only a part of it (Fig. 9–10). Because of that we consider that a real porosity of *T*. cf. *proschkinae* valves is smaller than 5 %.

The obtained result were compared with the data of Losic et al. [2007], who measured the porosity of two valves of centric diatoms, *T. eccentrica* and *Coscinodiscus sp*., using AFM (Table 1). Also De Stefano et al. [2008] estimated the porosity of the valve of *C. wailesii* using thresholding of SEM image, and obtained the value 0.57. As this data is very different from all previously obtained, and the authors did not provide a clear algorithm by which their data were obtained, we consider that this value is incorrect and do not include it into table.

## Discussion

The described method helps to study porosity of diatoms using SEM images where areolae and cribral pores are well visible. It is a good alternative to the studying of valve topography using AFM [Luis et al. 2017], because presented method does not require the cultivation of diatoms and gives the comparable results. Moreover presented method allows fulfilling retrospective analysis of species porosity from published images.

The described method is similar to the morphometric method of Sicko-Goad et al. [1977], who were estimated the volumes of the organelles of planktonic algae on electron microscope micrographs.

Manual selection of frustule (valve) components on an image is time consuming and appropriate for tiny diatoms with small number of areolae as those we consider. For a large diatoms with numerous areolae it is necessary to combine manual and automatic segmentation algorithms. Besides this it is possible to simulate the valve patterns and estimate the porosity of frustule components with the help of geometric models [Lyakh 2013].

The value of porosity is a component of the investigation of the functional morphology of a diatom frustule [Bukhtiyarova 2009, 2015]. It can be used as a quantitative character in diatoms taxonomy, and as a potential indicator of the influence of the environmental factors on microalgae.

The results obtained show that a very low valve area – less than 5 % of the total area – is sufficient for the providing a protoplast with nutrients and dissolved gases. It is interesting that this value is consistent with the relative perforation of land plants leaves, which is less than 3% [Lawson & Blatt 2014].

The correspondence between the porosity of valves of *M. vodyanitskiyi, T. eccentrica, Coscinodiscus* sp., and leaves of land plants allow to assume that such value of porosity is usual for many other diatom species. To verify this hypothesis we are planning to continue our researches.

## Acknowledgements

This work was supported by the scientific research theme of the Institute of Marine Biological Researchers RAS (No. AAAA-A18-118020790229-7). A part of the work was carried out in the Collective Instrumental Center “Ultramicroanalysis” within the framework of the basic project of the Ministry of Science and Higher Education of Russian Federation at the Limnological Institute of the Siberian Branch of RAS (No. 0345-2019-0001).

